# Single-cell analysis reveals the *KIT* D816V mutation in hematopoietic stem and progenitor cells in systemic mastocytosis

**DOI:** 10.1101/394304

**Authors:** Jennine Grootens, Johanna S. Ungerstedt, Maria Ekoff, Elin Rönnberg, Monika Klimkowska, Rose-Marie Amini, Michel Arock, Stina Söderlund, Mattias Mattsson, Gunnar Nilsson, Joakim S. Dahlin

**Author notes:** Corresponding authors at: Department of Medicine, Division of Immuology and Allergy, Karolinska Institutet, 17164 Stockholm, Sweden. Email addresses (G.N. Nilsson), (J.S. Dahlin).

## Abstract

**Background:** Systemic mastocytosis (SM) is a hematological disease characterized by organ infiltration by neoplastic mast cells. Almost all SM patients have a mutation in the gene encoding the tyrosine kinase receptor KIT causing a D816V substitution and autoactivation of the receptor. Mast cells and CD34^+^ hematopoietic progenitors can carry the mutation, however, in which progenitor cell subset the mutation arises is unknown. We aimed to investigate the distribution of the D816V mutation in single mast cells and single hematopoietic stem and progenitor cells.

**Methods:** Fluorescence-activated single-cell index sorting and D816V mutation assessment were applied to analyze mast cells and more than 10,000 CD34^+^ bone marrow progenitors across 10 hematopoietic progenitor subsets. In vitro assays verified cell-forming potential.

**Findings:** We found that in SM 60-99% of the mast cells harbored the D816V mutation. Despite increased frequencies of mast cells in SM patients compared with control subjects, the hematopoietic progenitor subset frequencies were comparable. Nevertheless, the mutation could be detected throughout the hematopoietic landscape of SM patients, from hematopoietic stem cells to more lineage-primed progenitors. In addition, we demonstrate that FcεRI+ bone marrow progenitors exhibit mast cell-forming potential, and we describe aberrant CD45RA expression on SM mast cells for the first time.

**Interpretation:** The *KIT* D816V mutation arises in early hematopoietic stem and progenitor cells and the mutation frequency is approaching 100% in mature mast cells, which express the aberrant marker CD45RA.

**Research in Context:** *Evidence before this study:* Systemic mastocytosis is a myeloid neoplasm associated with a mutation in the receptor KIT. The mutation is found in multiple mature cell lineages and can be detected in bulk-sorted CD34^+^ hematopoietic stem and progenitor cells.

*Added value of this study:* We use single-cell analysis to study the distribution of the common *KIT* mutation within the heterogenous CD34^+^ progenitor landscape and in mast cells of systemic mastocytosis patients. Isolation of progenitor subsets followed by cell culture identifies a population of progenitors in bone marrow with increased mast cell-forming potential. Furthermore, a novel marker for aberrant mast cells in systemic mastocytosis is described.

*Implications of all the available evidence:* Our study provides novel insights into the cellular origin and propagation of the common *KIT* mutation in systemic mastocytosis. The identification of a novel marker for aberrant mast cells shows potential to improve disease diagnosis.

## Introduction

Hematopoietic stem cells (HSCs) in the bone marrow give rise to blood cells and mast cells. Differentiating HSCs progress through a number of intermediate progenitors with multilineage-forming capacity before commitment to the mast cell lineage.^1,2^ The binding of stem cell factor (SCF) to its receptor, KIT, promotes the maturation and proliferation of mast cells.^3–6^ It is therefore not surprising that mutations in the *KIT* gene coincide with the mast cell-driven disease systemic mastocytosis (SM).^7^

SM is a hematologic neoplasm in which infiltrates of neoplastic mast cells occur in various tissues.^8,9^ The majority of SM patients carry a mutation in *KIT*, most commonly affecting codon 816.^10^ The *KIT* D816V mutation makes receptor signaling constitutively active, independent of binding to its ligand SCF. The detection of *KIT* D816V in either bone marrow or peripheral blood samples is one of the criteria for the clinical diagnosis of SM using a standardized qPCR assay.^11^ When the *KIT* D816V mutation is present in mast cells, it may also be detected in mature bone marrow or peripheral blood cells, such as basophils, eosinophils, neutrophils, and B and T lymphocytes, which depends on the patient.^12–17^ Furthermore, the precursors of erythroid and myeloid cells as well as CD34^+^ progenitors may carry the *KIT* D816V mutation.^12,18,19^ Taken together, these findings suggest that the mutation may arise in early hematopoietic stem and progenitor cells (HSPCs), rather than in a lineage-restricted precursor.

The CD34^+^ cell compartment is highly heterogeneous, spanning HSCs to late lineage-committed progenitors. In the present study, we delineated the cellular origin of the *KIT* D816V mutation in single bone marrow cells in SM by combining FACS index sorting of CD34^+^ HSPCs with a multiplex qPCR assay. The data revealed that the D816V mutation in *KIT* arises in early HSPCs. The mutation burden is low but variable in multipotent and lineage-restricted progenitor populations and increases in mature mast cells. Furthermore, the present study provides more insight into hematopoiesis in SM subjects and identifies high CD45RA expression in aberrant mast cells.

## Materials and Methods

### Patients and sample preparation

Bone marrow and occasional peripheral blood samples were collected from patients under evaluation for SM and from control subjects. The study was approved by the Stockholm and Uppsala Regional Ethics committee. Oral and written informed consent was obtained from each patient and control subject. The study was conducted in accordance with the World Medical Association Declaration of Helsinki. The patients included in our experimental studies were classified according to the World Health Organization criteria^20^ as having indolent SM (ISM, n = 19), aggressive SM (ASM, n = 3), SM with associated hematological neoplasm (SM-AHN, n = 3), or other malignancies, including myelodysplastic syndrome or myeloproliferative neoplasm (MDS or MPN, n = 2). Supplementary Table 1 shows the patient characteristics. For analytic purposes, ASM and SM-AHN patients were combined in an advanced SM group (AdvSM). The controls (Ctrl) included patients with cutaneous mastocytosis (CM, n = 2) and subjects without SM (n = 7). All samples were processed on the day of collection. Red blood cells were lysed with PharmLyse buffer (BD Biosciences, Franklin Lakes, NJ) and washed with PBS containing 2% fetal calf serum (FCS, Thermo Fisher Scientific, Waltham, MA).

### Cell lines

Two human mast cell lines, HMC-1.2 and ROSA^KIT WT^, with and without the *KIT* D816V mutation, respectively, were cultured as previously described.^21,22^

### Flow cytometry and cell sorting

Bone marrow cells were stained with fluorophore-conjugated monoclonal antibodies in Brilliant Stain Buffer (BD Biosciences, Franklin Lakes, NJ). The antibodies used for staining included CD10 (HI10a), CD14 (M5E2), CD34 (581), CD38 (Hb7), CD45RA (HI100), CD90 (5E10), CD117 (104D2), CD123 (6H6), CD133 (AC133), and FcεRIα (CRA-1) (all from BD Biosciences, Biolegend, San Diego, CA, or Miltenyi Biotec, Bergisch Gladbach, Germany). The live/dead cell marker DAPI (BD Bioscience) was used for the analysis of cultured cells. Cell sorting and analysis were performed using the FACSAria III and LSRFortessa systems (BD Biosciences). For cell culture experiments, cells were sorted into sterile-filtered PBS with 2% FCS following a two-step sorting protocol, using the yield followed by the purity precision mode. The analysis of 20-100 sorted cells verified the sorting purity. Single CD34^+^ cells were sorted into 2 μl of lysis buffer containing 20 μg/ml PCR-grade proteinase K (Thermo Fisher Scientific) in EB buffer (Qiagen, Hilden, Germany). These cells were sorted either immediately from the sample or following an enrichment sort. The single-cell precision mode ensured that only one cell was deposited per well. The plates were then centrifuged and stored at −80°C until analysis. The sorting data from single cells were saved using the index sort module of the software FACSDIVA software version 8 (BD Biosciences). Flow cytometry data analysis was performed using FlowJo software version 10 (TreeStar, Ashland, OR).

### Cell culture and colony-forming assays

Colony-forming potential was evaluated by plating 500 cells in 35 × 10 mm tissue culture dishes (Thermo Fisher Scientific) in duplicate. The cells were cultured in Methocult H4434, including SCF, IL-3, EPO, and GM-CSF (STEMCELL Technologies Inc., Vancouver, Canada). Colonies were counted after 12-13 days, distinguishing erythroid, granulocyte/monocyte, and mixed colonies following the manufacturer’s recommendations. The mast cell-forming potential was assessed via liquid culture as described previously.^23^ Briefly, sorted cells were cultured in medium with recombinant human (rh) IL-3 (10 ng/mL; PeproTech, Rocky Hill, NJ) and rhIL-6 (10 ng/mL; PeproTech). In some experiments, the medium was changed on day 5, and the cells were subsequently cultured with rhIL-6 (10 ng/mL) and SCF (100 ng/mL, r-metHuSCF; Swedish Orphan Biovitrum, Stockholm, Sweden), as indicated in the figure legends. Enzyme cytochemical staining of trypsin-like activity was used to assess tryptase expression.^24,25^ Cell morphology was evaluated by May-Grünwald Giemsa staining (MGG; Sigma-Aldrich, St. Louis, MO).

### Single-cell mutation analysis

Before the addition of lysis buffer, 96-well plates (Bio-Rad Laboratories, Hercules, CA) were UV-treated for 30 minutes using a UVT-B-AR UV Cabinet (Grant Instruments, Cambridge, UK). Tubes and pipettes were treated with the DNA AWAY Surface Decontaminant (Thermo Fisher Scientific). After sorting, the cells were lysed for 45 minutes at 55°C, followed by a 10-min heat-inactivating step at 95°C. Multiplex qPCR analysis was performed via a Taqman gene expression assay (Thermo Fisher Scientific) in lysed single cells using 96-well CFX Manager (Bio-Rad Laboratories). A mutation assay was designed based on a previously published method^11^ using Primer3 (v.0.4.0) which specifically detected *KIT* c.2447A>T (D816V) using the forward primer 5'-AGAGACTTGGCAGCCAGAAA-3' a reverse primer matching the mutation at the 3' position, 5'-TTAACCACATAATTAGAATCATTCTTGATGA-3' and the probe 5'-FAM-TCCTCCTTACTCATGGTCGGATCACA-BHQ1-3'. A control assay was designed to target a flanking exon and intron eight in GAPDH using the forward primer 5'-CTGACTTCAACAGCGACACC-3' the reverse primer 5'-AGAGTTGTCAGGGCCCTTTT-3' and the probe 5'-HEX-TCAAGCTCATTTCCTGGTATGTG-BHQ1-3' (all primers came from TAG Copenhagen, Denmark). The control and mutation assays were tested and optimized using the cell lines ROSA^KIT WT^ and HMC-1.2. The primer concentrations, optimized for single cells, were as follows: control assay 150 nM forward/reverse primers, 100 nM probe; mutation assay 100 nM forward/reverse primers, 67 nM probe. Each plate, containing 86 CD34^+^ cells, 5 mature mast cells, 3 empty wells, and 2 wells with bulk CD34^+^ cells following sorting, was subjected to the multiplex qPCR assay. Cells were excluded from analysis when amplification failed in the control assay, or when amplification of the control assay resulted in a CT value higher than 40. In the mutation assay, cells were regarded as unmutated when there was no amplification or when the CT value was higher than 45. Outliers from each plate were then detected using the ROUT outlier test in Prism software and removed from further analyses (Supplementary Fig. 6B). Plates and cells that passed the quality control procedure were linked with the index data and gated similarly to the whole sample for analysis.

### Statistical analysis

Statistical analysis was performed using Prism software (Version 6.0h, GraphPad Software, La Jolla, CA). The unpaired two-tailed Student’s t-test was employed when comparing two groups, and the unpaired one-way ANOVA with Tukey’s multiple comparison test was employed when comparing more than two groups. Statistical analysis of log-transformed data was performed for some experiments as indicated in the figure legends. Differences were considered significant when *P* < 0.05.

## Results

### Systemic mastocytosis patients have a normal CD34^+^ bone marrow stem and progenitor cell composition

To assess the composition of the hematopoietic cells in the bone marrow of SM patients, we designed a multicolor flow cytometry panel that identifies ten distinct HSPC subtypes and mature mast cells (Table 1; Fig. 1A-B). Colony formation assays confirmed the validity of the GMP, CMP, and MEP gating strategies in SM subjects (Supplementary Fig. 1A-C). The flow cytometry data provided a specific HSPC profile for 34 bone marrow samples: 9 control, 19 ISM and 6 AdvSM (Fig. 1C; Supplementary Table 1). The percentages of HSPCs in SM samples did not significantly differ from those in control samples (Fig. 1D and Supplementary Fig. 2A). In addition, the ISM and AdvSM samples exhibited percentages of total CD34^+^ cells similar to the control samples (Supplementary Fig. 2B). Thus, the overall hematopoiesis profile did not differ between the control and SM subjects.

**Table 1.** Immunophenotypes of hematopoietic stem and progenitor cells

| Short | Cell type | Gating strategy | References |
| --- | --- | --- | --- |
| HSC | Hematopoietic stem cell | CD14<sup>-</sup> CD34<sup>+</sup> CD38<sup>-</sup> CD45RA<sup>-</sup> CD90<sup>+</sup> | 26, 27 |
| MPP | Multipotent progenitor | CD14<sup>-</sup> CD34<sup>+</sup> CD38<sup>-</sup> CD45RA<sup>-</sup> CD90<sup>-</sup> | 26, 27 |
| CMP<sup>FcεRI<sup>-</sup></sup> | Common myeloid progenitor FcεRI<sup>-</sup> | CD14<sup>-</sup> CD34<sup>+</sup> CD38<sup>+</sup> CD45RA<sup>-</sup> CD10<sup>-</sup> CD123<sup>+</sup> FcεRI<sup>-</sup> | 28 * |
| CMP<sup>FcεRI<sup>+</sup></sup> | Common myeloid progenitor FcεRI<sup>+</sup> | CD14<sup>-</sup> CD34<sup>+</sup> CD38<sup>+</sup> CD45RA<sup>-</sup> CD10<sup>-</sup> CD123<sup>+</sup> FcεRI<sup>+</sup> CD117<sup>+</sup> | 28 * |
| MEP | Megakaryocyte-erythroid progenitor | CD14<sup>-</sup> CD34<sup>+</sup> CD38<sup>+</sup> CD45RA<sup>-</sup> CD10<sup>-</sup> CD123<sup>-</sup> | 28 |
| LMPP | Lymphoid-primed multipotent progenitor | CD14<sup>-</sup> CD34<sup>+</sup> CD38<sup>-</sup> CD45RA<sup>+</sup> | 29 |
| GMP | Granulocyte-monocyte progenitor | CD14<sup>-</sup> CD34<sup>+</sup> CD38<sup>+</sup> CD45RA<sup>+</sup> CD10<sup>-</sup> CD123<sup>+</sup> | 28 |
| CDP | Common dendritic progenitor | CD14<sup>-</sup> CD34<sup>+</sup> CD38<sup>+</sup> CD45RA<sup>+</sup> CD10<sup>-</sup> CD123<sup>hi</sup> | 30, 31 |
| MLP | Multilymphoid progenitor | CD14<sup>-</sup> CD34<sup>+</sup> CD38<sup>-</sup> CD45RA<sup>+</sup> CD10<sup>+</sup> | 32 |
| BNKP | B- and NK cell progenitor | CD14<sup>-</sup> CD34<sup>+</sup> CD38<sup>+</sup> CD45RA<sup>+</sup> CD10<sup>+</sup> | 32 |
| MC | Mast cell | CD14<sup>-</sup> CD34<sup>-</sup> CD117<sup>hi</sup> FcεRI<sup>+</sup> |  |

\*Refers to the original CMP population without the inclusion of the FcεRI marker.

**Fig. 1.**
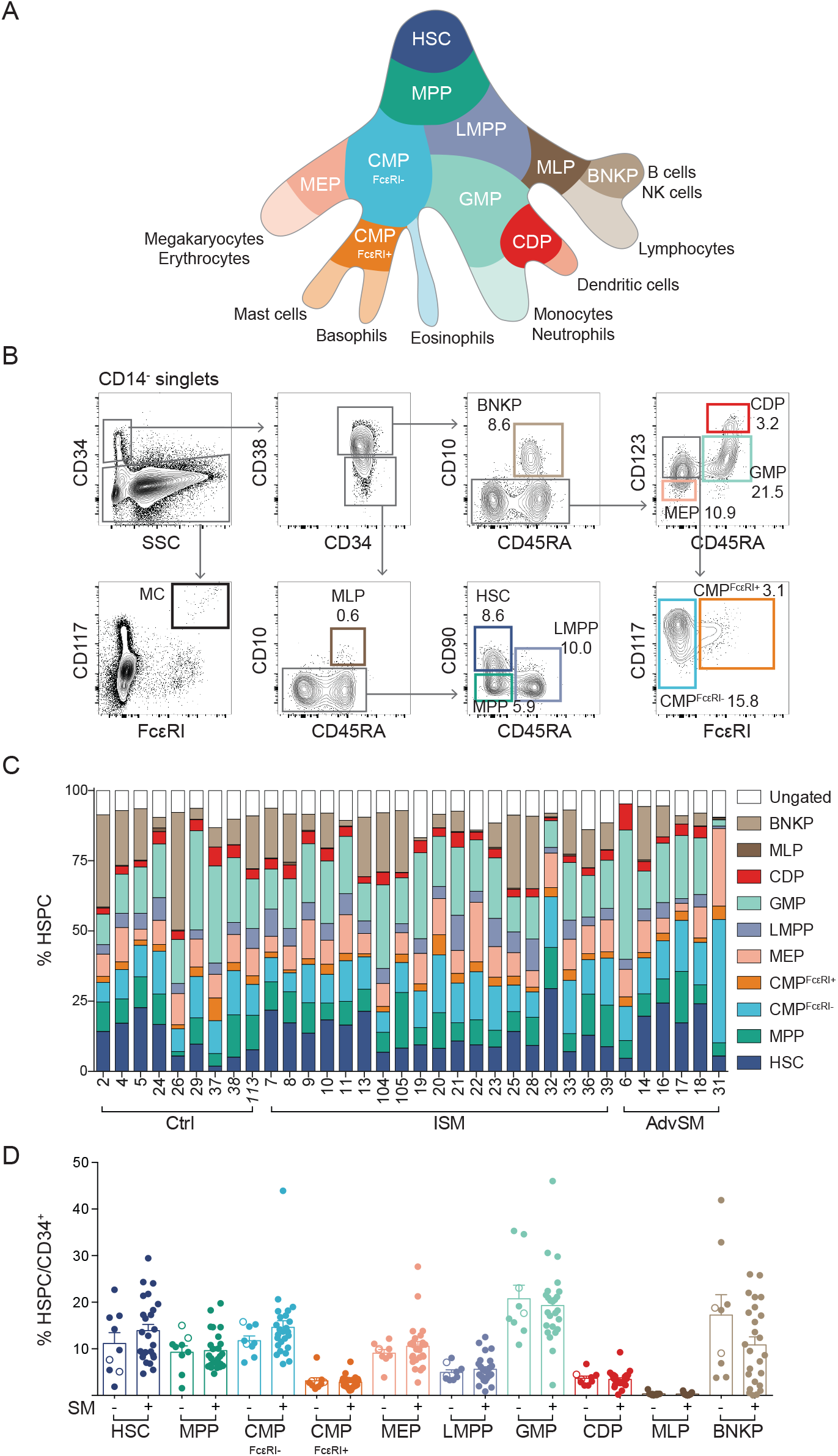
HSPC gating and profiling shows a normal course of hematopoiesis in patients with systemic mastocytosis. (A) A landscape model of the hematopoiesis. (B) Gating strategy for bone marrow HSPCs and mature mast cells, showing representative plots with population frequencies as percentages. (C) The hematopoietic progenitor profile of 34 bone marrow samples. Cutaneous mastocytosis samples are indicated in italics. (D) SM samples were compared with control samples without SM, including CM (open circles), and the mean expression + SEM of each HSPC is presented as a percentage of the total CD34^+^ cells. The unpaired *t*-test was used in panel D to compare the control and SM groups for each HSPC subset.

### The bone marrow CMP^FcεRI+^ fraction presents an enhanced mast cell-forming potential

Mast cell progenitors circulate in the blood as Lin^-^ CD34^hi^ CD117^int/hi^ FcεRI^+^ cells.^33^ These cells express CD123 and lack CD45RA,^23,33^ overlaying the CMP^FcεRI+^ gate in bone marrow (Supplementary Fig. 3A) that exhibit a distinct expression profile compared with mature mast cells (Supplementary Fig. 3B). We therefore analyzed the mast cell-forming potential of the bone marrow CMP^FcεRI+^ fraction by culturing the cells with IL-3 and IL-6. Flow cytometry analysis revealed that the CMP^FcεRI+^ cultures gave rise to a higher frequency of CD117^hi^ FcεRI^+^ mast cells than the CMPs^FcεRI-^ cells (Fig. 2A-C), despite overall lower CD117 expression (Supplementary Fig. 4A). May-Grünwald Giemsa and enzymatic tryptase staining showed that the CMP^FcεRI+^ progeny generated a high frequency of granulated tryptase-positive mast cells when cultured with SCF (Fig. 2D-E and Supplementary Fig. 5). Cultured GMPs did not give rise to any cells with a mast cell phenotype (Fig. 2A-E and Supplementary Fig. 5).

**Fig. 2.**
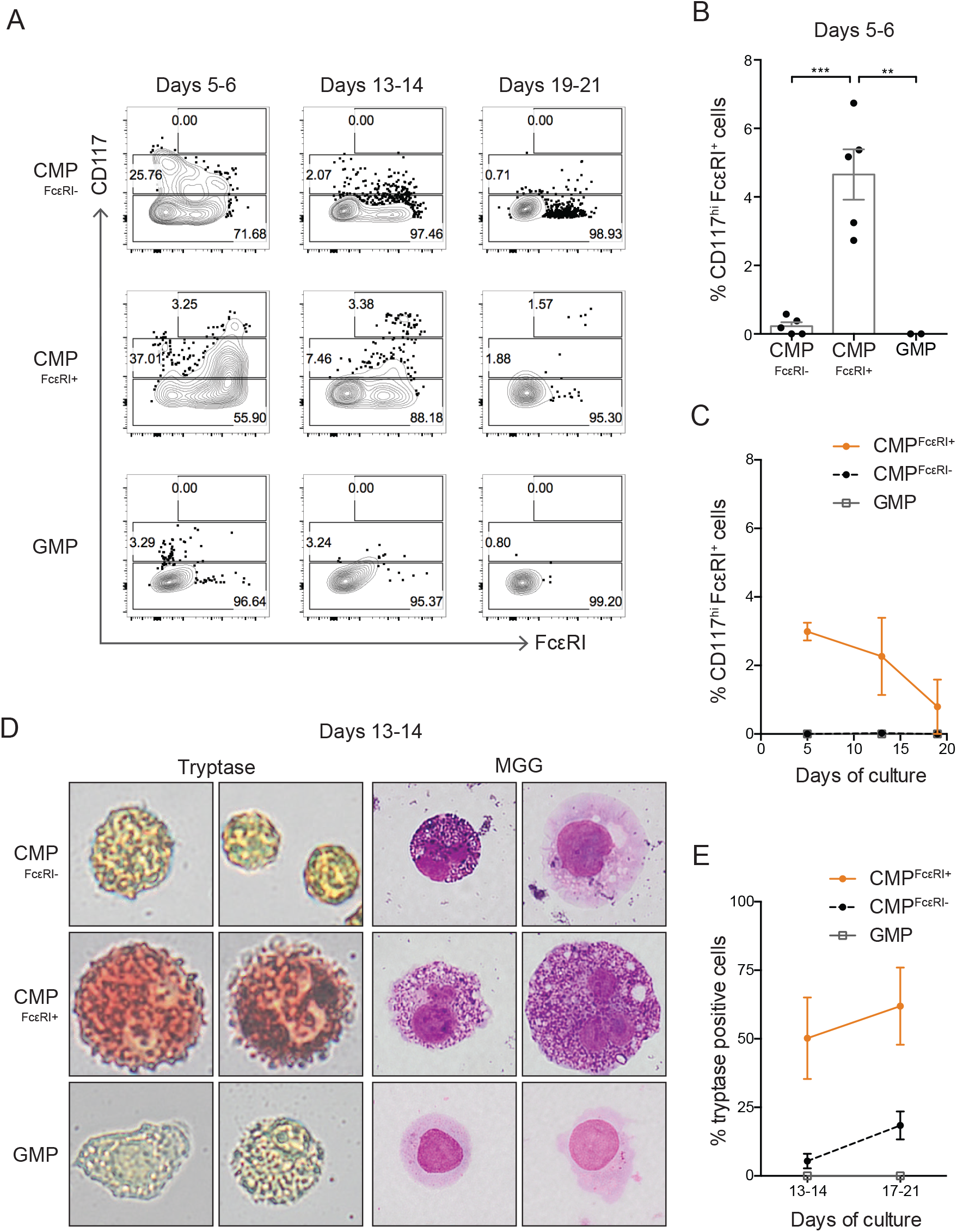
The CMP^FcεRI+^ fraction in the bone marrow has mast cell potential. (A-C) Flow cytometry analysis of of CMP^FcεRL·^, CMP^FcεRI+^ and GMP cells cultured with rhIL-3 and rhIL-6 up to 21 days. (A) Flow cytometry plots of representative cultures, showing the percentages of CD117^-^, CD117^+^, and CD117^hi^ cells in each plot. (B) Percentage of CD117^hi^ FcεRI^+^ cells at days 5-6. (C) Two of the samples in panel B were followed for up to 21 days under culture with rhIL-3 and rhIL-6. The percentage of CD117^hi^ cells was analyzed on days 5-6, 13-14, and 19-21. (D) Representative images of CMP^FcεRI-^, CMP^FcεRI+^ and GMP cells cultured with rhIL-3 and rhIL-6 for 5 days, followed by rhIL-6 and SCF until day 13-14, showing tryptase staining (red) and granules by May-Grünwald Giemsa (MGG) staining. (E) Quantification of tryptase-positive cells at days 13-14 and 17-21. CMP^FcεRL·^, CMP^FcεRI+^ cells were analyzed from Ctrl29, SM28, SM32, SM33, and MDS30. GMPs were analyzed from SM32 and SM33. The bars and lines in panel B, C and D represent the means ± SEM. One-way ANOVA with Tukey’s multiple comparison was used in panel B, \*\**P* <0.01, \*\*\**P* < 0.001. Images were captured using an Olympus XC10 camera (Olympus, Tokyo, Japan). The image width corresponds to 29 μm.

The emerging model of hematopoiesis proposes that MEPs, basophils, and eosinophils develop separately from GMPs following an asymmetric cell division at the MPP stage. This process can be traced by the segregation of CD133, which is expressed in GMPs but not in progenitors with erythromyeloid output.^34^ Indeed, we found that CMP^FcεRI+^ cells exhibited only low levels of CD133 staining, similar to erythromyeloid cells (Supplementary Fig. 4B).

### CD45RA is expressed on aberrant mast cells in systemic mastocytosis

The multicolor flow cytometry panel used for the gating of HSPCs was also employed to identify mast cells in bone marrow (Fig. 1B). May-Grünwald Giemsa and tryptase staining confirmed that CD14^-^ CD34^-^ CD117^hi^ FcεRI^hi^ cells constitute mast cells (Fig. 3A). SM samples presented higher percentages of mast cells than controls (Fig. 3B). Notably, mast cells from almost all SM samples exhibited aberrant CD45RA expression, as verified with fluorescence minus one control and internal control staining of CD34^+^ cells (Fig. 3C-D). The cutaneous (non-systemic) mastocytosis samples and MDS/MPN samples expressed low levels of CD45RA on bone marrow mast cells (Fig. 3C-D). In addition, mast cell frequencies were correlated with the clinical percentage of BMMC infiltrate and serum tryptase levels (Fig. 4A-C; Supplementary Table 2). Taken together, the results indicate that SM bone marrow samples contain more mast cells than control samples and these mast cells express high levels of the aberrant marker CD45RA.

**Fig. 3.**
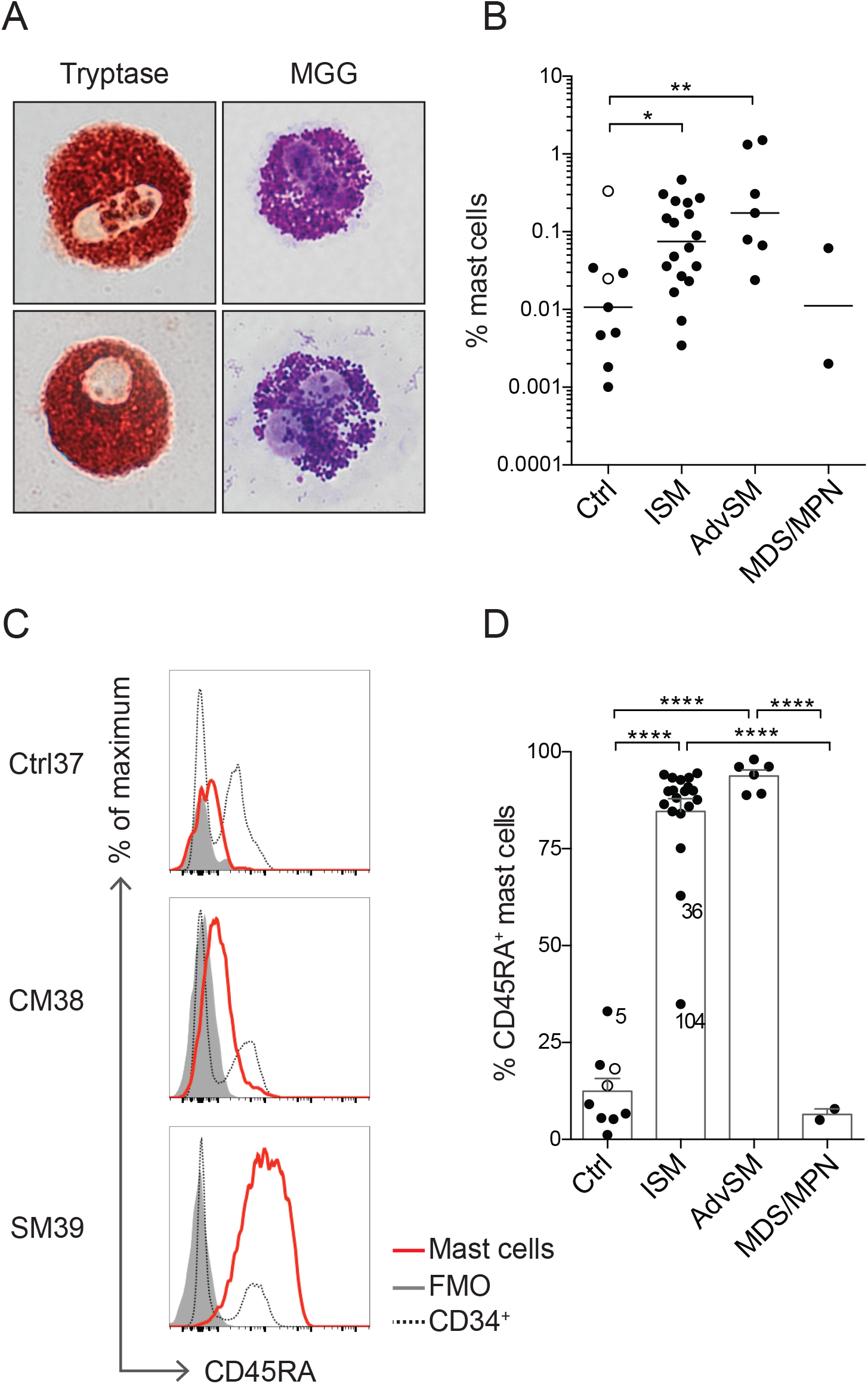
SM mast cells aberrantly express CD45RA. (A) Enzymatic staining demonstrates tryptase expression, and MGG staining reveals granules in SM mast cells. The images are representative of two experiments (samples SM32 and SM33). Images were captured using an Olympus XC10 camera (Olympus, Tokyo, Japan). The image width corresponds to 29 μm. (B) Mast cell frequencies in Ctrl, ISM, AdvSM, and MDS/MPN patients, showing the median of log-transformed values. (C) Histograms showing CD45RA expression in mast cells (red line), FMO controls (grey histogram) and CD34^+^ cells (dotted line). (D) Frequency of CD45RA-expressing mast cells, showing the mean and SEM. Control subjects included two subjects with CM (open circles). One-way ANOVA with Tukey’s multiple comparison was used in panels B and D, \**P* < 0.05, \*\**P* < 0.01, **** *P* < 0.0001.

**Fig. 4.**
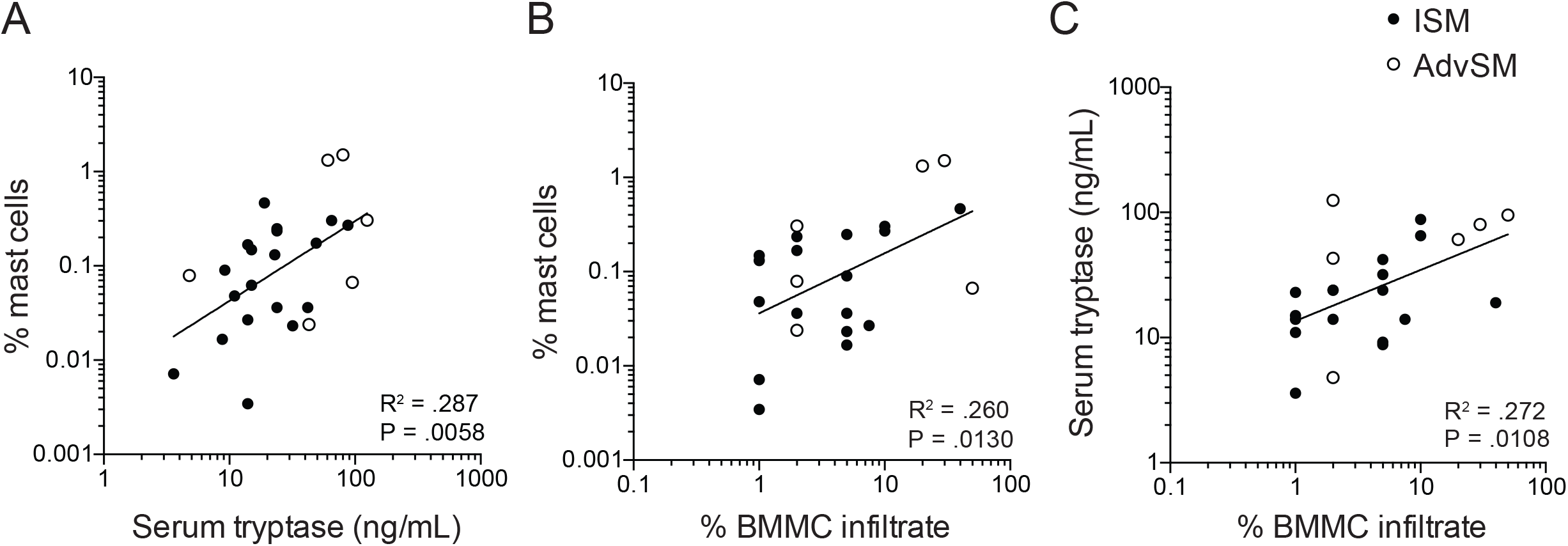
The percentage of mast cells in bone marrow correlates with clinical features. The percentage of mast cells was correlated with (A) serum tryptase levels and (B) the percentage of BMMC infiltrate in the biopsy section. (C) Serum tryptase levels were correlated with the percentage of BMMC infiltrate in the biopsy section. The samples include ISM (closed dots) and AdvSM (open dots). Correlation values and significance are indicated in the figure and in Supplementary Table 2. Linear regression was calculated based on log-transformed data: in panel A, n =25, and in panels B and C, n = 23. The percentage of BMMC infiltrate was not determined for two patients, accounting for the difference in numbers between panels. For analytical purposes, BMMC infiltrate presented as <1% or <5% in Supplementary Table 1 were set at 1% and 5% respectively.

### Single-cell index sorting reveals the KIT D816V mutation in early HSPCs in systemic mastocytosis patients

The cellular origin of the *KIT* D816V mutation in SM has not been established, and the relative abundance of this mutation in different CD34^+^ HSPC populations and mast cells has not been determined. We therefore developed a single-cell method to trace the *KIT* D816V mutation using two cell lines: HMC-1.2 cells carrying the mutation and ROSA cells carrying wild-type *KIT*. The cells were labeled with two different fluorochrome-labeled antibodies, mixed, and randomly FACS sorted with the index sort function. A multiplex qPCR assay was used to detect an irrelevant gene, for quality control purposes, and the *KIT* D816V mutation (Supplementary Fig. 6A-B). Among all single cells, more than 96% passed the quality control procedure (Supplementary Fig. 6C). Linking the results with the index data revealed that the mutation assay was negative for the wild-type ROSA cells. The assay detected the mutation in 98.1% of all HMC-1.2 cells. Thus, the assay did not detect false positive cells in the mutation assay and only a few false negatives, demonstrating the high sensitivity and specificity of the method (Supplementary Fig. 6D).

The high success rate with cell lines allowed us to use the assay for HSPCs and mature mast cells from patient samples. Single cells from 12 samples (6 ISM and 6 AdvSM) were analyzed for mutation assessment. Specifically, CD34^+^ HSPCs were index sorted from ten patients, whereas specific HSPC subsets were sorted from two patients. At the time of SM diagnosis, the presence of the *KIT* D816V mutation was confirmed in either peripheral blood or bone marrow (Supplementary Table 1). We analyzed 40-194 single mast cells and 351-1583 single CD34^+^ cells from each bone marrow sample, providing mutation data for 12,859 single cells in total (Fig. 5A; Supplementary Table 3). All SM samples carried the *KIT* D816V mutation in the mast cell compartment at a high frequency in ISM (mean, 80.9%) and AdvSM (mean, 93.2%) (Fig. 5B). Index data analysis of individual mast cells demonstrated that CD45RA expression was associated with presence of the *KIT* mutation (Fig. 5C). The percentage of mutated mast cells was correlated with the total percentage of mast cells in the sample and with the serum tryptase levels (Fig. 6A-B; Supplementary Table 2). The mutation rate in CD34^+^ HSPCs was less than 3% in all samples (Fig. 5D), except for one AdvSM sample with a high mutation burden of 37.0% (Supplementary Table 4). Nevertheless, the percentage of mutated CD34^+^ HSPCs was correlated with serum tryptase levels (Fig. 6C; Supplementary Table 2).

**Fig. 5.**
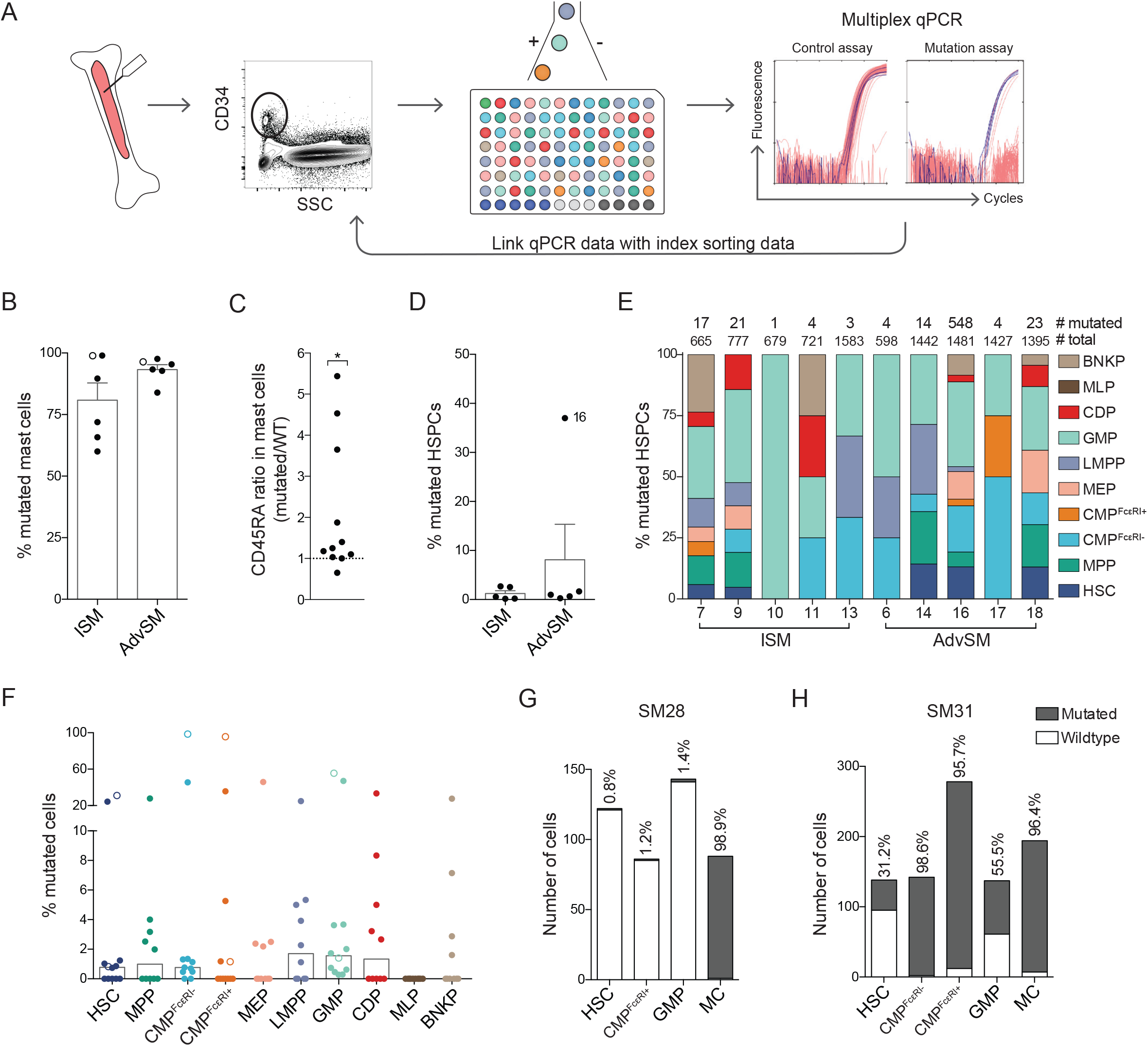
Mutation analysis of KIT D816V in single cells identifies mutated HSPCs and mast cells. (A) Method of single-cell mutation analysis using index sorting and multiplex qPCR. (B) The bars indicate the means + SEM percentage of mutated mast cells. (C) The ratio between the median CD45RA expression of mutated and wild type (WT) mast cells was determined for each patient. Each data point represents one patient. The numbers of cells analyzed from each patient are presented in Supplementary Table 3. Eleven patients are shown, as index data was not available from 1 of the 12 patients. The two-tailed Wilcoxon signed rank test determined whether the ratio was different from 1, indicated with a dotted line. \**P* < 0.05. (D) The bars indicate the means + SEM percentage of mutated HSPCs. (E) The cellular distribution of the mutation in 10 different HSPCs, showing the total numbers of mutated and sorted cells for each sample that passed the quality control procedure. (F) The bars indicate the median percentage of mutated cells per HSPC. (G-H) Mutation frequencies of selectively isolated HSPCs following a 2-step sorting protocol in two samples (SM28, ISM; and SM31, AdvSM). These two samples are indicated with open circles in panels B and F.

**Fig. 6.**
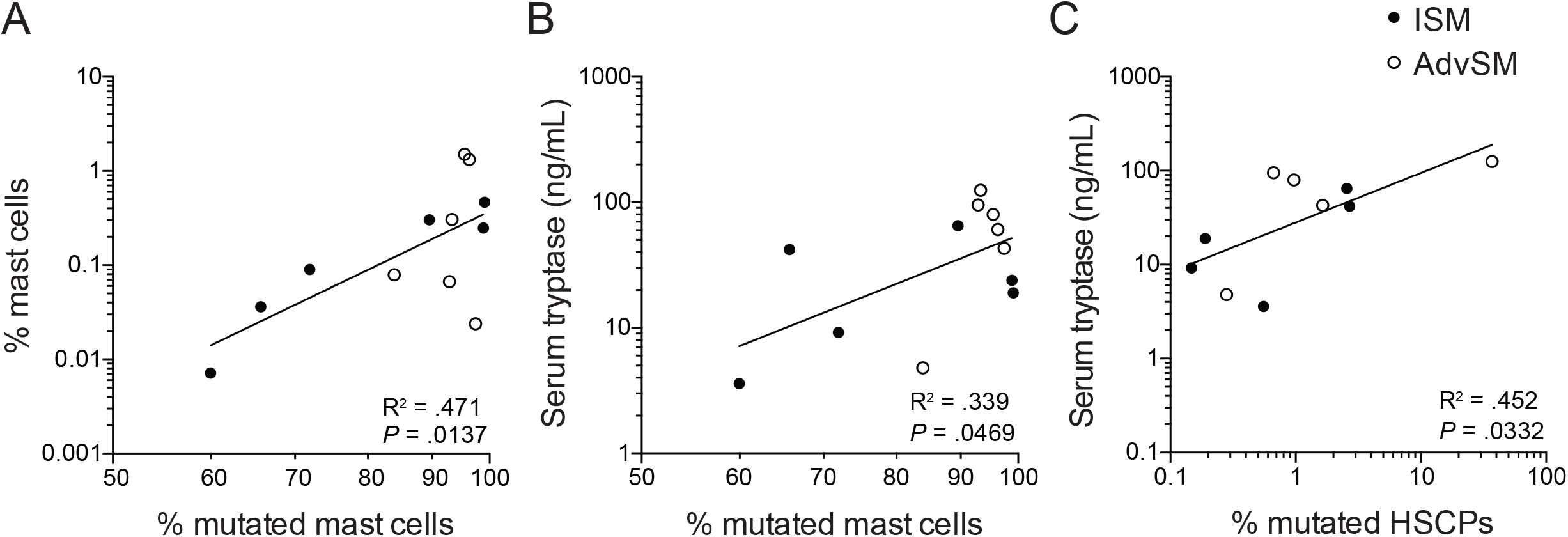
The mutation rate in mast cells and HSPCs correlates with clinical features. (A-C) Linear regression of the percentage of mutated mast cells, showing significant correlations with (A) the percentage of mast cells and (B) serum tryptase levels. (C) The percentage of mutated HSPCs was correlated with serum tryptase levels. The samples included ISM (closed dots) and AdvSM (open dots). Correlation and significance are indicated in the figure and in Supplementary Table 2. Linear regression was calculated based on log-transformed data in panels A-C, n = 12 and D, n = 10.

Coupling the index data with the mutation analysis of 10 bone marrow samples revealed that the *KIT* D816V mutation appeared throughout the hematopoietic landscape (Fig. 5E; Supplementary Table 3-4). Specifically, different HSPCs presented similar mutation rates in all samples when the median percentage of mutation was compared between different HSPCs (Fig. 5F). Overall, the method of index sorting provided a representative set of single cells from the whole-bone-marrow aspirates (Supplementary Fig. 7A). However, the CMP^FcεRI+^ population with an enriched mast cell potential was rare. To determine the mutation frequency in the CMP^FcεRI+^ fraction, we selectively sorted 86 and 278 single CMPs^FcεRI+^ from 2 SM samples. One patient exhibited less than 2% mutated CMPs^FcεRI+^, HSCs and GMPs, whereas the fraction of mutated mast cells was 98.9% (Fig. 5G; Supplementary Table 4). The second patient presented 95.7% mutated CMPs^FcεRI+^ and 96.4 *%* mutated mast cells (Fig. 5H; Supplementary Table 4). Peripheral blood MCPs (CD14^-^ CD34^+^ CD117^+^ FcεRI^+^) showed mutation rates comparable to that of bone marrow CMPs^FcεRI+^ in one patient (Supplementary Fig. 7B), but the mutation could not be detected in the MCPs from 4 other SM samples, likely due to the low mutation frequency and the small number of cells analyzed due to the rarity of the cells (14-23 cells analyzed per sample). Taken together, we detected the *KIT* D816V mutation throughout the hematopoietic landscape, starting with the HSCs.

## Discussion

The cellular origin of the aberrant clone in SM is still under debate. Here, we show that SM patients exhibit an increased frequency of mast cells in bone marrow. Almost all these mast cells carry the *KIT* D816V mutation. Despite normal HSPC frequencies in bone marrow, the *KIT* D816V mutation can be traced all the way back to the CD34^+^ CD38^-^ CD10^−^ CD45RA^-^ CD90^+^ HSC fraction.

The *KIT* D816V mutation causes receptor autoactivation, which signals survival, proliferation, and differentiation, likely explaining the competitive growth advantage of the aberrant mast cell clone and the high frequency of mutated mast cells. However, the frequency of mutated HSPCs was low in most patients. This observation is consistent with the overall normal HSPC composition of SM patients. There are two main potential explanations for the low D816V mutation burden observed in HSPCs relative to mast cells: HSPCs rapidly differentiate into mast cells when the mutation occurs, or mutated HSPCs do not present a strong growth advantage over non-mutated HSPCs. In fact, we have previously shown that KIT signaling is dispensable for mast cell progenitor development in vitro and in vivo, and that other stimuli can induce cell proliferation.^23^ However, the aberrant progenitor clone may in some cases proliferate to reach a high frequency, as demonstrated in two patients with AdvSM who presented more than 20% mutated cells in the HSC fraction. Advanced forms of SM have previously been associated with detection of the *KIT* mutation in bulk-sorted CD34^+^ progenitors and various mature cell lineages, whereas in indolent disease, the mutation is mainly restricted to mast cells.^13^

It is of particular interest to study the mutation burden in progenitors with mast cell-forming potential. We have previously reported that CD34^+^ CD117^int/hi^ FcεRI^+^ cells in peripheral blood are committed or nearly committed MCPs.^33^ Considering the CD34^+^ CD117^+^ FcεRI^+^ phenotype of CMPs^FcεRI+^ cells, these cells might be expected to present a similar mast cell-forming capacity as to blood MCPs. In the current study, we showed that CMPs^FcεRI+^ gave rise to a higher frequency of mast cells than CMPs^FcεRI-^ and GMPs. However, the mast cell frequency in cultured CMPs^FcεRI+^ was less than 10% and a substantial fraction were found to be CD117^-/lo^ FcεRI^+^ cells after culture, consistent with a basophil-like phenotype. Thus, CMPs^FcεRI+^ exhibit a considerable mast cell-forming capacity but also include progenitors with non-mast cell output. Basophils mature in bone marrow, whereas mast cells mature in peripheral tissues, which likely explains why bone marrow CMPs^FcεRI+^ form CD117^-^ FcεRI^+^ basophil-like cells, whereas CD34^+^CD117^int/hi^ FcεRI^+^ blood progenitors do not. The high frequency of CD117^-^ FcεRI^+^ basophil-like cells that develop from CMPs^FcεRI+^ is also in agreement with the reduced CD117 expression of CMPs^FcεRI+^ compared with that of CMPs^FcεRI-^. CD117 is upregulated upon mast cell maturation, and blood mast cell progenitors express intermediate to high levels of CD117. CMPs^FcεRI+^ express CD117 at low levels overall, even though there are individual CMPs^FcεRI+^ cells that express high CD117 levels, which might correspond to the cells with mast cell-forming capacity. Taken together, our results indicate that the low *KIT* mutation prevalence in the CMP^FcεRI+^ fraction is likely explained by the capacity of these cells’ to form not only mast cells but also CD117^-^ FcεRI^+^ basophil-like cells.

In SM, mast cells are known to aberrantly express multiple surface proteins, among CD2 and CD25 are currently used for diagnosis.^35^ These markers are expressed independently of the SM subtype. Other markers, such as CD30 and CD52, are associated with AdvSM and mast cell leukemia.^36,37^ In the present study, we demonstrate that CD45RA is a novel marker for aberrant mast cells in SM. Remarkably, bone marrow mast cells from two patients with cutaneous mastocytosis expressed low CD45RA levels, suggesting that CD45RA is a marker for systemic mast cell disease. The increase in the CD45RA isoform of CD45 likely explains the high levels of CD45 observed in systemic mastocytosis patients.^38^–^41^ Taken together, our results indicate that high CD45RA expression should be considered a novel marker for diagnosis of SM.

CD45RA is present on aberrant mast cells in SM, but at what stage during mast cell differentiation is the marker upregulated? CMPs^FcεRI+^, which lack CD45RA, present mast cell-forming potential in SM patients. In contrast, we show that cells in the classic CD45RA^+^ GMP gate lack mast cell potential in SM patients. An alternative explanation is that the mast cell progenitors in SM fall into the immature CD38^-^ gate. In fact, examples of rare mutated CD117^int/hi^ FcεRI^int/hi^ cells in the CD45RA^+^ LMPP gate were seen (Supplementary Fig. 7C). It is also tempting to speculate that CD45RA is upregulated once aberrant mast cells are mature.

The CD34^+^ HSPC population is highly heterogeneous and a single-cell resolution is therefore necessary to delineate the cellular distribution of the mutation. Here, we used single-cell index sorting to track cellular identity when performing mutation analysis. Notably, the mutation was detected throughout differentiation, from HSCs to the MEP, GMP, and LMPP fractions, depending on the patient. The detection of the mutation in early HSPCs explains previous findings of the mutation being present in bulk-sorted mature cell lineages of some patients.

Multiplex TaqMan qPCR assays detected the *KIT* mutation in index-sorted progenitors in the present investigation. The qPCR method was originally developed for sensitive mutation detection in DNA extracted from bone marrow or blood cells. GAPDH primers and probes were added to the assay to confirm the presence or absence of a sorted cell. This method allowed the detection of mutated cells with close to 100% efficiency, with no observed false positives. The single-cell index sorting and DNA mutation analysis approach can similarly be applied to other mutations and hematologic diseases to reveal the origin of aberrant clones. One limitation of the method is the number of cells that can be analyzed, meaning that mutated cells in very rare subpopulations might go undetected. However, more than 10,000 HSPCs were analyzed in the present investigation, and single-cell index sorting coupled with qPCR allowed unbiased and prospective analysis of mutated HSPC populations in SM. Whole-genome amplification and nested PCR methods for amplifying and sequencing DNA mutations from single cells are associated with the phenomenon known as allelic dropout, which is the amplification of only a single allele, leading to an overestimation of the frequency of non-mutated cells. Mutational profiling of single colonies, as performed by Jawhar *et al*.^18^ avoids this problem. However, such analysis can be performed only on progenitors that form sufficiently large colonies. Mast cell progenitors present a poor proliferative capacity, and this technique was therefore not applied in the present investigation.

In summary, we have explored the composition and mutation profile of CD34^+^ HSPCs from SM patients. The results show that the *KIT* mutation can be found in cells throughout the hematopoietic landscape with increased burden in the mast cell lineage, supporting the notion that the aberrant clone may arise in the hematopoietic stem cell compartment. Furthermore, the mutated mast cells express CD45RA, a potential novel clinical biomarker for aberrant mast cells in SM.

## Acknowledgements

We wish to thank Kerstin Hamberg Levedahl for help with the patients and Petter Woll for helpful discussions. Stem cell factor was a kind gift from SOBI, Stockholm, Sweden.

## Funding Sources

This study was supported by the Swedish Research Council, the Swedish Cancer Society, the Stockholm Cancer Society (Radiumhemmets Forskningsfonder), Magnus Bergvall’s Foundation, Tore Nilsson’s Foundation for Medical Research, the Karolinska Institutet, and through the regional agreement on medical training and clinical research (ALF) between Stockholm County Council and the Karolinska Institutet, and ALF-funding from Uppsala University Hospital.

## Declaration of Interest

The authors declare no competing interests

## Author Contributions

Conceptualization and Methodology, J.G., G.N. and J.S.D.; Investigation, J.G., M.E. and E.H.; Formal analysis, J.G.; Visualization and Validation, J.G. and J.S.D.; Resources, J.S.U., M.K., R.A., S.S., M.M. and M.A.; Writing – original draft, J.G. and J.S.D.; Writing – review & editing, all authors; Funding acquisition and Supervision, J.U., G.N. and J.S.D.

