## Supplementary data for "Single-cell analysis reveals the *KIT* D816V mutation in hematopoietic stem and progenitor cells in systemic mastocytosis"

#### Supplementary figure legends

**Supplementary Fig. 1**
**Differentiation potential of CD38$^+$ HSPCs in SM.** (A) Gating strategy used for sorting CMPs, GMPs, and MEPs. (B) Distribution of granulocyte/monocyte, erythroid, or mixed colonies, counted on days 12-13. (C) Representative colonies. CMPs, GMPs, and MEPs were sorted from samples SM21, SM22, and SM23; whereas CMPs and GMPs were sorted from SM20. Images were captured using a Zeiss AxioVert 200 Inverted microscope (Zeiss, Oberkochen Germany).

**Supplementary Fig. 2**
**SM subjects are similar to the controls but differ from MDS/MPN subjects.** (A) Percentage of HSPCs per total CD34$^+$ cells comparing the control, ISM, AdvSM groups with the MDS/MPN subjects. (B) The percentage of CD34$^+$ cells in controls, ISM and AdvSM and MDS/MPN subjects. Means and SEM are shown. Unpaired one-way ANOVA with Tukeys comparisons were used in panel A and B, $^{**}P < 0.01$, $^{***}P < 0.001$.

**Supplementary Fig. 3**
**Expression profile of mast cells and CMP$^{Fc\varepsilon RI+}$ cells in PB and BM.**
(A) Representative gating of MCPs in PB showing CD123, CD45RA and CD34 expression. (B) Backgating analysis of CMPs$^{Fc\varepsilon RI+}$ and mature mast cells in BM of a representative subject. CMPs$^{Fc\varepsilon RI+}$ and mature mast cells are overlaid in the bottom panels.

**Supplementary Fig. 4**
**CD133 and CD117-expression on HSPCs.** (A) CD117 expression on CMP$^{Fc\varepsilon RI+}$ cells was lower than on CMP$^{Fc\varepsilon RI-}$ cells in 6 randomly selected samples, showing the median fluorescent intensity of CD117 expression. (B) CD133 expression on HSPCs from samples Ctrl113, SM19, and SM36, showing the median fluorescent intensity for the stained sample minus the FMO control. Expression levels were normalized within each sample by assigning the sum of the expression of all populations to as 100%.

**Supplementary Fig 5.**
**Staining for granularity and tryptase in cultured CMP$^{Fc\varepsilon RI-}$, CMP$^{Fc\varepsilon RI+}$ and GMP cells**.
Representative images of cultured cells from SM29 and SM33 on days 17-19. Images were captured using an Olympus XC10 camera (Olympus, Tokyo, Japan). The image width corresponds to 29 µm.

**Supplementary Fig. 6**
**The single-cell mutation assay shows high specificity and sensitivity in the mast cell lines HMC-1.2 and ROSA$^{KIT\ WT}$.** (A) Method of single-cell analysis in the mast cell lines ROSA$^{KIT\ WT}$ and HMC-1.2 (*KIT* D816V). Cells were labeled separately for CD117 and sorted using the index sort option. Each cell was analyzed for a control assay and mutation assay. Mutation data were linked to the index data. Bulk cell sorting was used as a control for the multiplex qPCR. (B) Flow diagram of the quality control performed for the single cells after qPCR. (C) Numbers and percentages of cells that showed amplification in the control assay for the wild-type and mutated cells. (D) Numbers and percentages of cells that showed amplification in the mutation assay.

**Supplementary Fig. 7**
**Details of single sorted HSPCs in bone marrow and peripheral blood.** (A) All single sorted cells identified by using the index-sorting data provided for each patient as a percentage of the total number of sorted cells. (B) Numbers and percentages of mutated cells in all HSPC populations including peripheral blood mast cell progenitors (PB MCP) in patient SM16. (C) FcεRI and CD117 expression on LMPPs in SM14.

#### Supplementary table legends

**Supplementary Table 1.** Patient characteristics.
**Supplementary Table 2.** Relations between clinical and experimental data.
**Supplementary Table 3.** Numbers of *KIT*-mutated and wildtype *KIT* hematopoietic stem and progenitor cells and mast cells.
**Supplementary Table 4.** Percentage of mutation per hematopoietic stem and progenitor cells and mast cells.

Supplementary Fig. 1

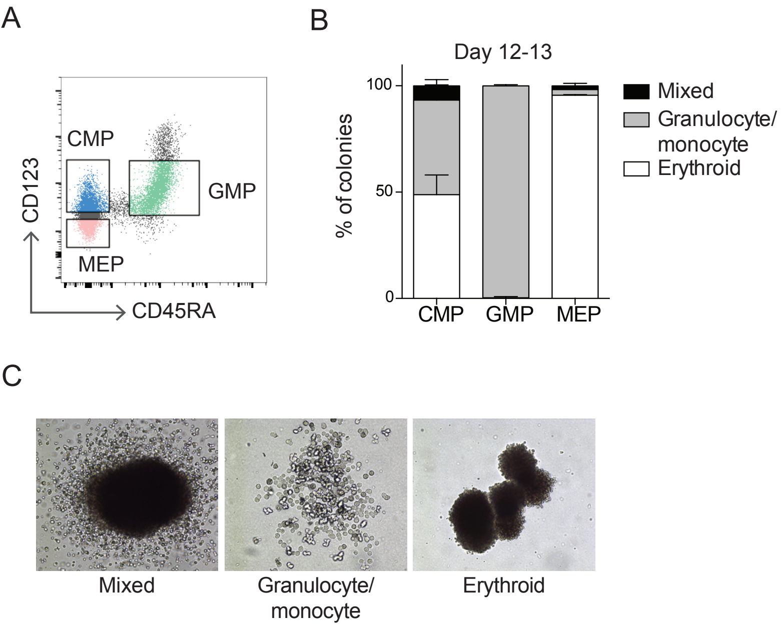

### Supplementary Fig. 2

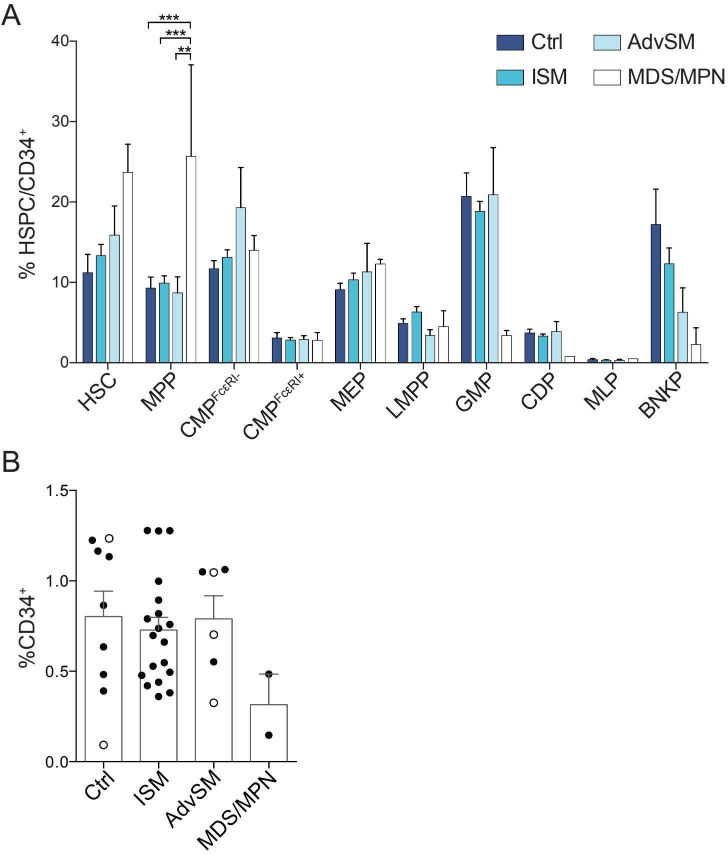

### Supplementary Fig. 3

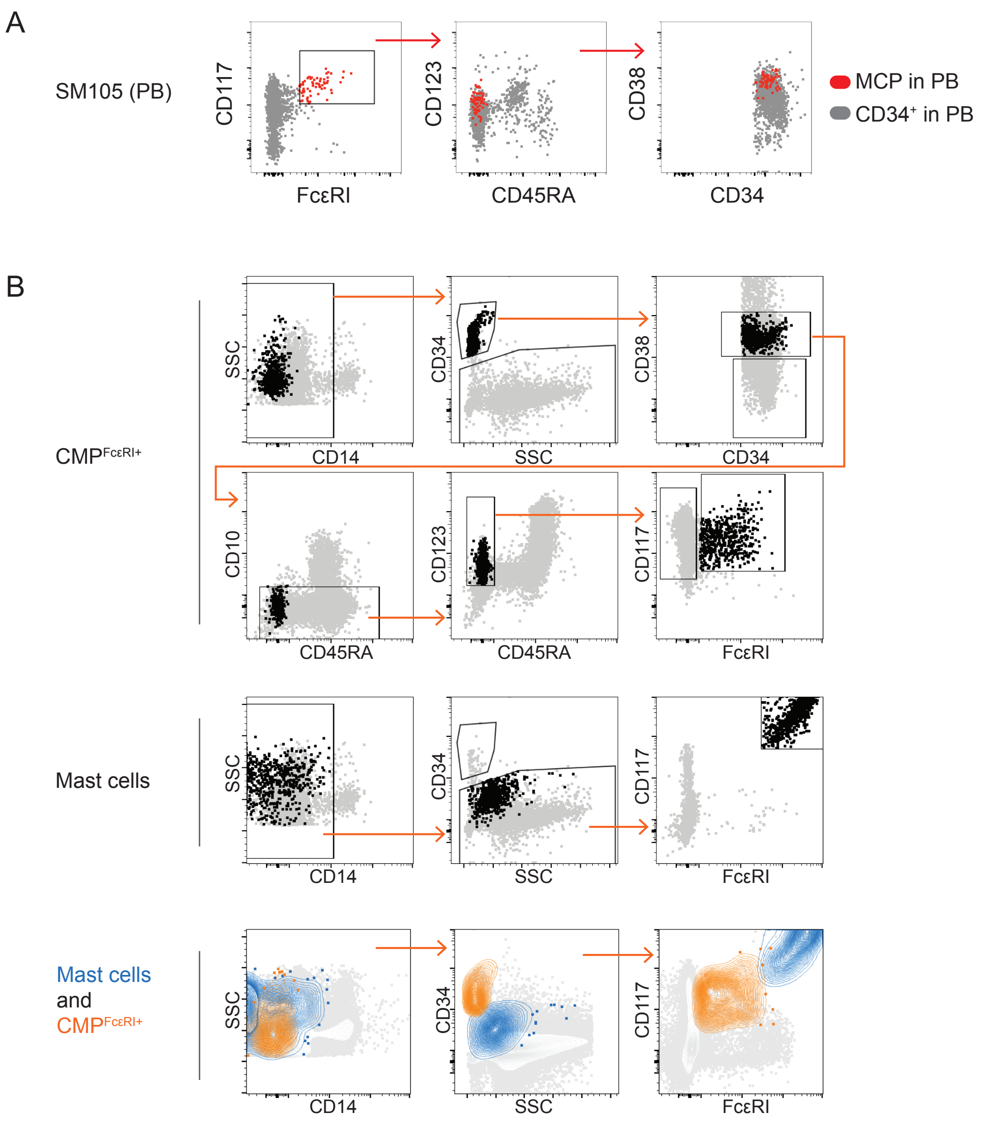

Supplementary Fig. 4

A

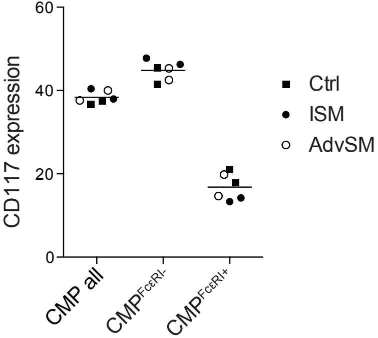


B

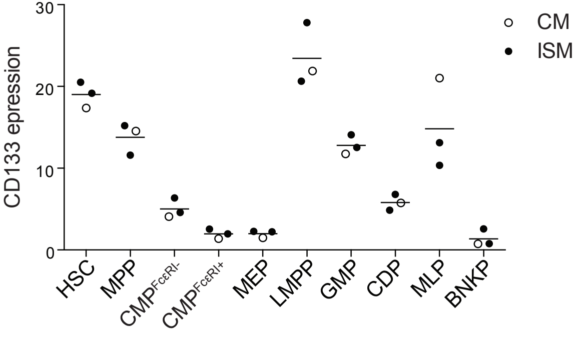

### Supplementary Fig. 5

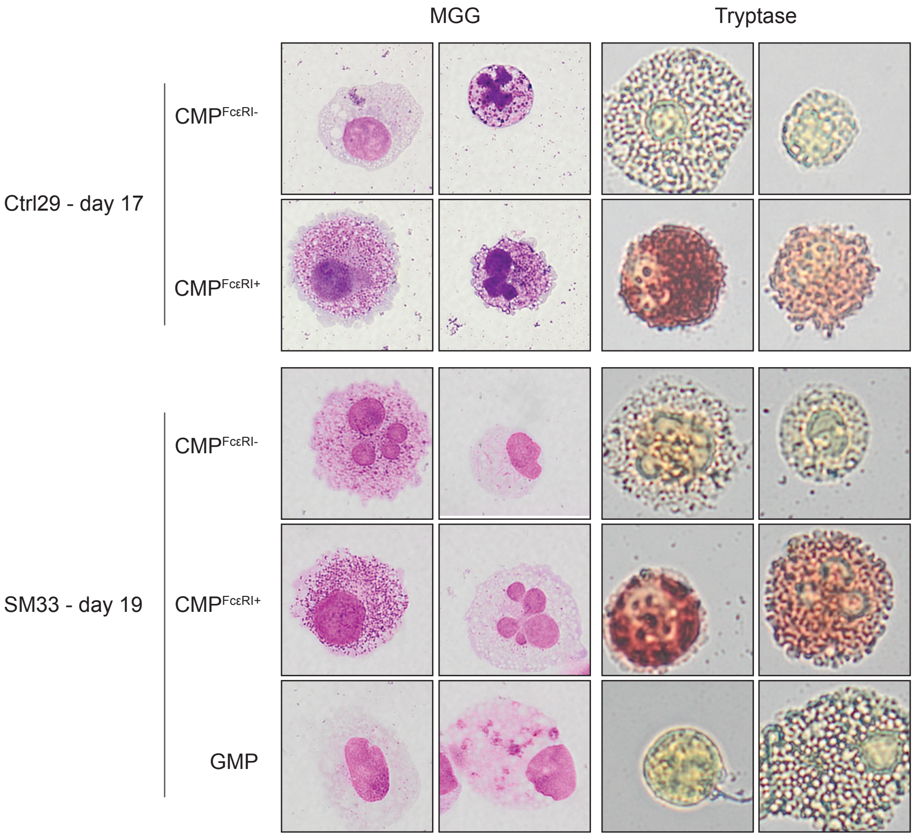

Supplementary Fig. 6

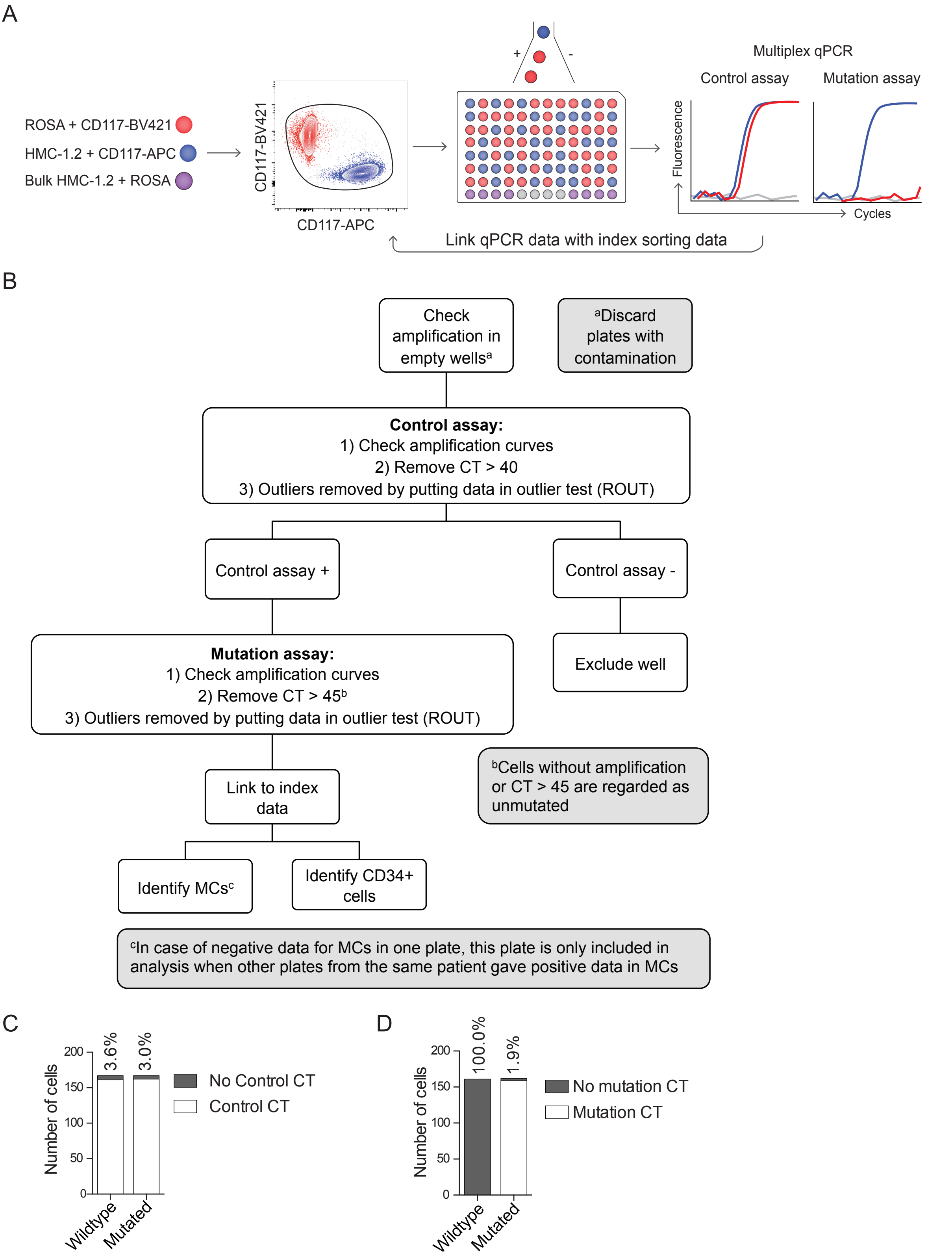

Supplementary Fig. 7

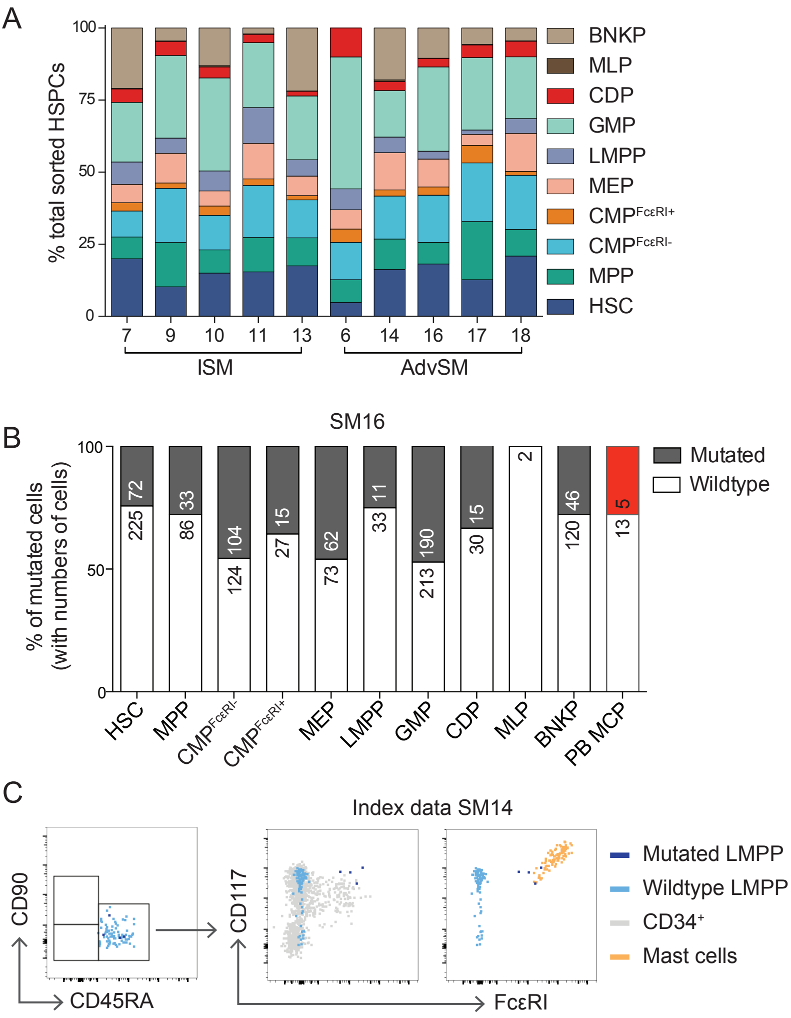

**Supplementary Table 1. Patient characteristics**.

| Patient | Gender | Age | WHO classification | Major criterion | BMMC infiltrate, % | Tryptase, ng/mL | D816V, PB | D816V, BM | Atypical MC morphology | CD2/ CD25 | UP |
|---|---|---|---|---|---|---|---|---|---|---|---|
| Ctrl02 | F | 29 | No SM | *NA* | *NA* | *NA* | *NA* | *NA* | *NA* | *NA* | *NA* |
| Ctrl04 | F | 73 | No SM | *NA* | *NA* | *NA* | *NA* | *NA* | *NA* | *NA* | *NA* |
| Ctrl05 | F | 35 | No SM | *NA* | *NA* | *NA* | *NA* | *NA* | *NA* | *NA* | *NA* |
| Ctrl24 | F | 42 | No SM | *NA* | *NA* | *NA* | *NA* | *NA* | *NA* | *NA* | *NA* |
| Ctrl26 | M | 37 | No SM † | *NA* | *NA* | *NA* | *NA* | *NA* | *NA* | *NA* | *NA* |
| Ctrl29 | M | 32 | No SM | *NA* | *NA* | *NA* | *NA* | *NA* | *NA* | *NA* | *NA* |
| Ctrl37 | F | 45 | No SM | *NA* | *NA* | *NA* | *NA* | *NA* | *NA* | *NA* | *NA* |
| Ctrl38 | M | 53 | CM * | *NA* | *NA* | *NA* | *NA* | *NA* | *NA* | *NA* | *NA* |
| Ctrl113 | M | 21 | CM * | *NA* | *NA* | *NA* | *NA* | *NA* | *NA* | *NA* | *NA* |
| SM07 | F | 53 | ISM | + | 10 | 65 | + | + | + | + | + |
| SM08 | F | 49 | ISM | - | <1 | 14 | + | + | + | + | + |
| SM09 | F | 65 | ISM | + | 5 | 42 | + | *ND* | - | + | + |
| SM10 | M | 26 | ISM | - | <5 | 9.2 | - | + | + | + | + |
| SM11 | F | 31 | ISM | - | <1 | 3.6 | + | + | + | + | + |
| SM13 | F | 65 | ISM | + | 40 | 19 | + | + | + | + | + |
| SM104 | F | 52 | ISM | - | <1 | 11 | - | + | - ‡ | + | - |
| SM105 | F | 70 | ISM | + | 2 | 24 | + | + | + | + | + |
| SM19 | F | 86 | ISM | - | <1 | 15 | + | + | + | + | + |
| SM20 | M | 58 | ISM | - | <1 | 23 | *ND* | + | + | + | + |
| SM21 | M | 46 | ISM | - | <5 | 8.8 | + | + | + | + | + |
| SM22 | M | 77 | ISM | + | 2 | 24 | *ND* | + | *ND* | + | - |
| SM23 | F | 45 | ISM | + | 5 | 32 | + | + | + | + | + |
| SM25 | F | 60 | ISM | + | 2 | 14 | *ND* | + | + | + | - |
| SM28 | F | 56 | ISM | + | 5 | 24 | *ND* | + | + | + | + |
| SM32 | F | 73 | ISM § | + | 10 | 88 | + | + | + | + | + |
| SM33 | M | 25 | ISM \|\| | *ND* | *ND* | 15 | + | + | *ND* | + | + |
| SM36 | F | 46 | ISM | - | 7.5 | 14 | + | + | + | + | + |
| SM39 | F | 65 | ISM | - | *ND* | 49 | + | + | - | + | + |
| SM06 | M | 85 | ASM ¶ | + | 50 | 95 | + | + | - | + | *ND* |
| SM14 | M | 44 | ASM | + | 30 | 80 | + | + | - | + | + |
| SM16 | M | 46 | ASM | + | 2 | 125 | *ND* | + | + | + | + |
| SM17 | F | 69 | SM-AHN # | + | 2 | 4.8 | *ND* | + | + | + | - |
| SM18 | M | 67 | SM-AHN # | + | 2 | 43 | *ND* | + | + | + | - |
| SM31 | F | 69 | SM-AHN ** | + | 20 | 61 | + †† | + †† | + | + | *ND* |
| MDS12 | F | 72 | MPN | - | *ND* | 18 | - | - | + | + | *ND* |
| MDS30 | M | 78 | MDS | - | *ND* | 17 | - | - | - | + | - |

BMMC, bone marrow mast cell; *NA*, not applicable; *ND*, not determined; CM, cutaneous mastocytosis; UP, Urticaria pigmentosa; PB, peripheral blood; BM, bone marrow; * Patient has cutaneous mastocytosis; † Patient has severe kidney failure; ‡ Second biopsy showed atypical morphology; § Small CLL clone in bone marrow, no peripheral lymphocytosis; || Bone marrow biopsy not available; ¶ ASM-AHN suspected; # SM-MPN; ** SM-MDS-MPN; †† High mutation burden

**Supplementary Table 2. Relations between clinical and experimental data.**

| | % Mutated mast cells | % Mutated HSPCs | % BMMC infiltrate | Serum tryptase | Age |
|---|---|---|---|---|---|
| % Mast cells | $R^2$ = .471; *P* = .0137; n = 12 | $R^2$ = .016; *P* = .7265; n = 10 | $R^2$ = .260; *P* = .0130; n = 23 | $R^2$ = .287; *P* = .0058; n = 25 | $R^2$ = .068; *P* = .2095; n = 25 |
| % Mutated mast cells | | $R^2$ = .029; *P* = .6356; n = 10 | $R^2$ = .254; *P* = .0948; n = 12 | $R^2$ = .339; *P* = .0469; n = 12 | $R^2$ = .294; *P* = .0688; n = 12 |
| % Mutated HSPCs | | | $R^2$ = .071; *P* = .4560; n = 10 | $R^2$ = .452; *P* = .0332; n = 10 | $R^2$ = .013; *P* = .7510; n = 10 |
| % BMMC infiltrate | | | | $R^2$ = .272; *P* = .0108; n = 23 | $R^2$ = .016; *P* = .5716; n = 23 |
| Serum tryptase | | | | | $R^2$ = .139; *P* = .0664; n = 25 |

**Supplementary Table 3. Numbers of *KIT*-mutated and wildtype *KIT* hematopoietic stem and progenitor cells and mast cells.**

| | SM7 | | SM9 | | SM10 | | SM11 | | SM13 | | SM6 | | SM14 | | SM16 | | SM17 | | SM18 | | SM28 | | SM31 | |
|---|---|---|---|---|---|---|---|---|---|---|---|---|---|---|---|---|---|---|---|---|---|---|---|---|
| | M | WT | M | WT | M | WT | M | WT | M | WT | M | WT | M | WT | M | WT | M | WT | M | WT | M | WT | M | WT |
| HSC | 1 | 134 | 1 | 80 | 0 | 102 | 0 | 112 | 0 | 278 | 0 | 28 | 2 | 232 | 72 | 225 | 0 | 182 | 3 | 291 | 1 | 121 | 43 | 95 |
| MPP | 2 | 48 | 3 | 116 | 0 | 55 | 0 | 86 | 0 | 154 | 0 | 50 | 3 | 148 | 33 | 86 | 0 | 288 | 4 | 122 | *ND* | *ND* | *ND* | *ND* |
| $CMP^{Fc\varepsilon RI-}$ | 0 | 61 | 2 | 146 | 0 | 81 | 1 | 129 | 1 | 207 | 1 | 75 | 1 | 215 | 104 | 124 | 2 | 288 | 3 | 261 | *ND* | *ND* | 140 | 2 |
| $CMP^{Fc\varepsilon RI+}$ | 1 | 18 | 0 | 15 | 0 | 22 | 0 | 17 | 0 | 23 | 0 | 30 | 0 | 32 | 15 | 27 | 1 | 84 | 0 | 19 | 1 | 85 | 266 | 12 |
| MEP | 1 | 41 | 2 | 78 | 0 | 36 | 0 | 89 | 0 | 107 | 0 | 38 | 0 | 187 | 62 | 73 | 0 | 55 | 4 | 179 | *ND* | *ND* | *ND* | *ND* |
| LMPP | 2 | 49 | 2 | 38 | 0 | 47 | 0 | 90 | 1 | 88 | 1 | 43 | 4 | 71 | 11 | 33 | 0 | 22 | 0 | 72 | *ND* | *ND* | *ND* | *ND* |
| GMP | 5 | 131 | 8 | 213 | 1 | 217 | 1 | 161 | 1 | 349 | 2 | 268 | 4 | 227 | 190 | 213 | 1 | 357 | 6 | 292 | 2 | 141 | 76 | 61 |
| CDP | 1 | 30 | 3 | 33 | 0 | 26 | 1 | 19 | 0 | 26 | 0 | 62 | 0 | 46 | 15 | 30 | 0 | 63 | 2 | 73 | *ND* | *ND* | *ND* | *ND* |
| MLP | 0 | 1 | 0 | 1 | 0 | 3 | 0 | 1 | 0 | 1 | 0 | 0 | 0 | 8 | 0 | 2 | 0 | 2 | 0 | 2 | *ND* | *ND* | *ND* | *ND* |
| B-NK | 4 | 135 | 0 | 36 | 0 | 89 | 1 | 13 | 0 | 347 | 0 | 0 | 0 | 262 | 46 | 120 | 0 | 82 | 1 | 61 | *ND* | *ND* | *ND* | *ND* |
| Mast cells | 60 | 7 | 25 | 13 | 41 | 16 | 24 | 16 | 97 | 1 | 103 | 8 | 83 | 4 | 83 | 6 | 78 | 15 | 81 | 2 | 87 | 1 | 187 | 7 |

M, number of cells with the *KIT* D816V mutation; WT, number of cells with wildtype *KIT*; *ND*, not determined

**Supplementary Table 4. Percentage of mutation per hematopoietic stem and progenitor cells and mast cells.**

| | SM7 | SM9 | SM10 | SM11 | SM13 | SM6 | SM14 | SM16 | SM17 | SM18 | SM28 | SM31 |
|---|---|---|---|---|---|---|---|---|---|---|---|---|
| HSC | 0.74 | 1.23 | 0.00 | 0.00 | 0.00 | 0.00 | 0.85 | 24.24 | 0.00 | 1.02 | 0.82 | 31.16 |
| MPP | 4.00 | 2.52 | 0.00 | 0.00 | 0.00 | 0.00 | 1.99 | 27.73 | 0.00 | 3.17 | *ND* | *ND* |
| $CMP^{Fc\varepsilon RI-}$ | 0.00 | 1.35 | 0.00 | 0.77 | 0.48 | 1.32 | 0.46 | 45.61 | 0.69 | 1.14 | *ND* | 98.59 |
| $CMP^{Fc\varepsilon RI+}$ | 5.26 | 0.00 | 0.00 | 0.00 | 0.00 | 0.00 | 0.00 | 35.71 | 1.18 | 0.00 | 1.16 | 95.68 |
| MEP | 2.38 | 2.50 | 0.00 | 0.00 | 0.00 | 0.00 | 0.00 | 45.93 | 0.00 | 2.19 | *ND* | *ND* |
| LMPP | 3.92 | 5.00 | 0.00 | 0.00 | 1.12 | 2.27 | 5.33 | 25.00 | 0.00 | 0.00 | *ND* | *ND* |
| GMP | 3.68 | 3.62 | 0.46 | 0.62 | 0.29 | 0.74 | 1.73 | 47.15 | 0.28 | 2.01 | 1.40 | 55.47 |
| CDP | 3.23 | 8.33 | 0.00 | 5.00 | 0.00 | 0.00 | 0.00 | 33.33 | 0.00 | 2.67 | *ND* | *ND* |
| MLP | 0.00 | 0.00 | 0.00 | 0.00 | 0.00 | *ND* | 0.00 | 0.00 | 0.00 | 0.00 | *ND* | *ND* |
| B-NK | 2.88 | 0.00 | 0.00 | 7.14 | 0.00 | *ND* | 0.00 | 27.71 | 0.00 | 1.61 | *ND* | *ND* |
| Total $CD34^{+}$ | 2.56 | 2.70 | 0.15 | 0.55 | 0.19 | 0.67 | 0.97 | 37.00 | 0.28 | 1.65 | *ND* | *ND* |
| Mast cells | 89.55 | 65.79 | 71.93 | 60.00 | 98.98 | 92.79 | 95.40 | 93.26 | 83.87 | 97.59 | 98.86 | 96.39 |

*ND*, not determined
